# A note on the history, morphology, and distribution ecology of the boreal felt lichen, *Erioderma pedicellatum* (Hue) P.M. Jørg. in the northern hemisphere

**DOI:** 10.64898/2026.10.22.760986

**Authors:** André Arsenault, Patricia Baines

## Abstract

The complex history of discovery, population declines, extinctions, and rediscoveries of the boreal felt lichen in the northern hemisphere is examined here for the first time in 124 years following its detection in Canada. We also present evidence of the presence of two distinct morphological forms of the boreal felt lichen. The southern form is in Atlantic Canada and is characterized by a darker, hairless exciple, shorter spores, and a P+ pruina. The northern form occurs in Norway, Alaska, Kamchatka, Quebec, Labrador and northwestern Newfoundland. Although the morphology of the northern form seems identical across North America, Europe, and Asia, the community ecology patterns for the lichen in Quebec’s Manicouagan region and western Labrador deviate from the other subpopulations. New genomic analyses will be required to untangle the population structure of the boreal felt lichen. Since the last IUCN assessment, the number of mature individuals has increased 42-fold, and its geographic distribution has expanded to include western north America, Asia, and the boreal mainland of Canada. This significant increase in population, combined with extensive un-surveyed suitable habitat, and the metapopulation dynamics of this lichen, will make future assessments challenging and will require more field research.

## Introduction

The boreal felt lichen, *Erioderma pedicellatum* (Hue) P.M. Jørg. is an arboreal cyanolichen in the *Pannariacae* family, mostly associated with conifer trees in the wet forests of the northern hemisphere (Ahti and Jørgensen 1971). It has been an emblem of global lichen conservation for over four decades (Clayden 1997). The presumed disappearance of this lichen from its type localities in New Brunswick, Canada (Hue 1911) and in Sweden (Ahlner 1944, 1948), and severe declines of known occurrences from clearcutting of old-growth forests in Canada (COSEWIC 2002), and in Sweden and Norway (Tønsberg et al. 1996) has led to several conservation concerns and actions.

These have included a designation of a critically endangered status on the Red List of Lichenized Fungi of the World by the International Committee for the Conservation of Lichens, Salzburg, Austria, 1996 (Scheidegger 1998) and a letter written from 94 members of the International Association of Lichenology representing 24 countries to the Honourable Premier of Newfoundland and Labrador, Brian Tobin, requesting him to take action to protect this lichen (Clayden 1997), and an official designation as critically endangered by the IUCN (Scheidegger 2003). The Committee on the Status of Endangered Wildlife in Canada (COSEWIC) assessed the status of the boreal felt lichen as endangered for the Atlantic population located in the Maritime provinces, and special concern for the boreal population located on the Island of Newfoundland (Cosewic 2002, 2014). In Europe, it was presumed extirpated in 1990 (Jørgensen 1990) but was found again in 1994 (Holien et a. 1995) and then presumed extinct again in 2003 (Scheidegger 2003) but was re-discovered the same year (Reiso and Hofton 2006).

The boreal felt lichen, *Erioderma pedicellatum* (Hue) P.M. Jørg. has a rich history involving near extinctions in Europe (Holien et al. 1995, Clayden 1997), local declines in Eastern Canada (Goudie et al. 2011, Cameron and Toms 2016), re-discoveries in Norway (Reiso and Hofton 2006), and new discoveries in the boreal regions of North America (Nelson et al. 2009, Anderson et al. 2022, Arsenault 2022, this paper) and Asia (Stepanchikova and Himelbrandt 2012) (Table 1).

**Table 1.** A summary of significant timelines associated with the distribution ecology of the boreal felt lichen, *Erioderma pedicellatum* (Hue) P.M. Jørg., syn, *Pannaria pedicellate Hue*, *Erioderma boreale* Ahlner. Sources of information are given below.

| Year | Location | Activity and pertinent voucher specimens |
| --- | --- | --- |
| 1902 | New Brunswick, Canada<sup>1</sup> | First collection of the southern form in Atlantic Canada on Passamaquoddy Bay, Campobello Island by William Gilson Farlow, not described until 1911. |
| 1911 | New Brunswick, Canada<sup>2</sup> | First species description of the southern form by Father Auguste Marie Hue, published as *Pannaria pedicellata* Hue Voucher (NHM-London- BM001097302 |
| 1938 | Nord-Trøndelag, Norway<sup>3</sup> | First collection of northern form in Europe by Sten Ahlner Voucher S-L70552 |
| 1939-1947 | Varmland, Sweden<sup>4</sup> | First collections of northern form in Sweden and a new collection from Norway by Sten Ahlner Vouchers S-L70551 & S-L70553 |
| 1944 | Scandinavia<sup>5</sup> | First description of the northern form as *Erioderma boreale* Ahlner by Sten Ahlner. |
| 1944 | Probably Alaska<sup>6</sup> | Presumed new collection of northern form in western North America by Sk. Harris and H. M. Raup but cannot verify location. It is reported from British Columbia but is not consistent with ecology of the area. Most |
|  |  | likely from Alaska. Voucher MIN-Lichens 941306 |
| 1948 | Scandinavia<sup>7</sup> | *Erioderma boreale* Ahlner (northern form) officially described in Sten Ahlner's Ph.D. dissertation. Voucher (NHM-London-BM001 097301) |
| 1956 | Newfoundland, Canada<sup>8</sup> | First collection of the southern form in Newfoundland by Ted Ahti but not identified until 1971. |
| 1971 | Newfoundland, Canada<sup>9</sup> | Identification of the Newfoundland southern form 1956 sample as *Erioderma boreale*. Voucher CANL 36037 |
| 1972 | New Brunswick, Canada<sup>10</sup> | Synonymization of *P. pedicellata* and *E. boreale* to, *E. pedicellatum* by Per Magnus Jørgensen. This combined the northern and southern form as one taxon. |
| 1978 | Costa Rica<sup>11</sup> | Mis-identified specimens. Vouchers: MIN-Lichens 941416, 941415, 941613 |
| 1980 | Newfoundland, Canada<sup>12</sup> | New Finds of southern form by Wolfgang Maas in different regions of Newfoundland. Voucher BG-L82418 |
| 1990 | Scandinavia<sup>13</sup> | Per Magnus Jørgensen declares the northern form of *E. pedicellatum* extinct in Scandinavia. |
| 1994 | Nord-Trøndelag, Norway<sup>14</sup> | Re-discovery of presumed extinct northern form in Scandinavia. Voucher TRH:L:4935/1 |
| 2001 | Quebec, Canada<sup>15</sup> | Northern form collected but not identified until 2021. |
| 2003 | Atlantic Canada<sup>16</sup> | Assessment by COSEWIC of the southern form of the Atlantic population as endangered and boreal population as special concern. |
| 2003 | Global<sup>17</sup> | IUCN assessment of global population as Critically Endangered A2c+4c |
| 2003 | Norway<sup>18</sup> | New subpopulation of northern form found in Rendalen Norway. |
| 2007 | Alaska, USA<sup>19</sup> | First official find of northern form in western North America. |
| 2009 | Kamchtaka, Russia<sup>20</sup> | First find of the northern form in Asia. |
| 2013 | Alaska, USA<sup>21</sup> | Population assessment of northern form published for the USA. |
| 2014 | Eastern Canada<sup>22</sup> | Re-assessment by COSEWIC of outhern form for the Atlantic population as Endangered C1+2a(i) and boreal population as Special Concern |
| 2019 | Kamchtaka, Russia<sup>23</sup> | Population assessment for the northern form published for Russia. |
| 2021 | Norway<sup>24</sup> | Population estimate reported for the northern form in Norway. |
| 2021 | Quebec, Canada<sup>25,26</sup> | First reported finds of northern form in Quebec. Voucher CDFN AFEL2021-2557 |
| 2021 | Labrador, Canada<sup>27</sup> | First find of northern form in Labrador. Voucher CDFN AFEL2021-2553 |
| 2025 | Gros Morne National Park.<sup>28</sup> | First find of northern form in Newfoundland Voucher CDFN AFEL2025X1 |

<sup>1</sup>(Pfister 2016), <sup>2</sup>(Hue 1911), <sup>3,4</sup>(Ahlner 1948), <sup>5</sup>Ahlner, S. (1944), presented the new species of the lichen *E. boreale*, which he found in Värmland and in N. Tröndelag at the Botanical Section meeting in Uppsala on May 9 1944 and reported in (Selling 1945), and in (Ahlner 1948), <sup>6</sup>GBIF (Toby Spribille & Daniel Stanton personal communication), <sup>7</sup>(Ahlner 1948), <sup>8</sup>CNHAL(collected by Ahti 1956, North-west shore of Long Pond), <sup>9</sup>Ahti and Jørgensen 1971, <sup>10</sup>Jørgensen 1972, <sup>11</sup>GBIF(Daniel Stanton, Personal communication), <sup>12</sup>GBIF( Maas 1980, many locations in Newfoundland), <sup>13</sup> (Jørgensen 1990), <sup>14</sup>GBIF(collection by Gaarder 1994, N. Tröndelag), <sup>15</sup>(Anderson et al. 2022), <sup>16</sup>(COSEWIC 2002), <sup>17</sup>(Scheidegger 2003) <sup>18</sup>(Reiso & Hofton 2006), <sup>19</sup>(Nelson et al. 2009), <sup>20</sup>(Stepanchikova and Himelbrant 2012), <sup>21</sup>(Stenh et al. 2013), <sup>22</sup>(COSEWIC 2014), <sup>23</sup> (Tagirdzhanova et al. 2019), <sup>24</sup> (Nilsson et al. 2022), <sup>25</sup>(Anderson et al. 2022), <sup>26</sup>(Arsenault 2022), <sup>27</sup>(This paper) <sup>28</sup>(This paper) .

The lichen was first collected in 1902 on Campobello Island, New Brunswick, Canada by Botanist William Gilson Farlow (Hue 1911; Jørgensen 1972; Pfister 2016). It remained un-reported until 1911 when lichenologist, Father Auguste-Marie Hue described it as a new lichen, *Pannaria pedicellata Hue*. (Hue 1911). It was later discovered in Norway in 1938 and in Sweden in 1939 by Botanist Sten Ahlner and officially presented as a new taxon, *Erioderma boreale* to the Swedish Botanical Association in 1944 (Selling 1945) and formally described as *Eriodema boreale* Ahlner. in a published Doctoral dissertation by Ahlner (1948) unaware that the same taxon had been previously published under a different name (Jørgensen 1972). Botanist, Per Magnus Jørgensen who specializes in the *Pannariacae* family (Jørgensen 2000) came upon the original description of *Pannaria pedicellata* and specimens from New Brunswick at the Natural History Museum of London and synonymized it with *E. boreale* as *E. pedicellatum* Hue P.M. Jørg. One early collection from western North America included a 1944 record by Harris and Raup from the Alaska highway. It was reported from British Columbia near summit Lake and although the date and Alaska highway reference matched lichen collections made by these lichenologists, the area near summit Lake, BC, does not represent suitable habitat for the boreal felt lichen (Toby Spribille and Trevor Goward, personal communication). It should be noted that there is a Summit Lake in Alaska in a similar region to where the boreal felt lichen has been found. It is therefore possible that this was the first collection of this lichen in the USA. In 1959, Ted Ahti collected the boreal felt lichen on the island of Newfoundland on the Avalon Peninsula. It was not identified and reported until 15 years later as *E. boreale* (Ahti and Jørgensen 1971). In 1978, it was reported from Costa Rica in 1978, but this turned out to be a misidentification (Daniel Stanton, personal communication). In the 1980’s Wolfgang Maas found this lichen in Nova Scotia and many other locations on the Island of Newfoundland (Cosewic 2002). In 2001 it was collected from Quebec’s North Shore east of the Manic 5 reservoir but was not identified and reported until twenty years later (Anderson et al. 2022). In 2003 a new subpopulation was found in Norway (Reiso and Hofton 2006) with over 1500 thalli on 6 spruce trees (Kjell Arne Solhaug, personal communication). The boreal felt lichen had major range extensions with new discoveries in Alaska in 2007 and Kamchatka in 2009. Further studies in these regions claimed they had the largest populations of the boreal felt lichen with inferred number of thalli inferred or approximated at one hundred thousand each (Stehn et al. 2013; Tagirdzhanova et al. 2019). In 2021, I (first author) discovered 4 new subpopulations of the boreal felt lichen in Quebec’s Manicouagan and western Labrador regions (Arsenault 2022) in addition to one collected twenty years prior in Quebec and reported in (Anderson et al. 2022). The latest find was by Parks Canada staff skiing in Gros Morne National Park in the spring of 2025 that had been trained on rare lichen surveys by myself and the land guardians of Miawpukek First Nation. This was a notable find because the thalli looked different and were only on branches of white spruce despite the presence of balsam fir in the same forest. It also suggested that in some forests detection of the boreal felt lichen could be impaired because in this case they were often above 3 metres from the ground.

When I (first author) started working on the ecology of the boreal felt lichen in 2010, I noticed some differences in morphology between the European pictures of this lichen and specimens I observed in the forests on the island of Newfoundland. As Jørgensen synonymized *E. boreale* and *P. pedicellata* to *E. pedicellatum*, he did notice some differences including a slightly more bulbous shape of the apothecia and more pronounced brown colour of the thalli of Canadian specimens. Genetic analyses of the mycobiont (Yetman 2007) and of the photobiont (Cornejo and Sheidegger 2016, Cornejo et al. 2016) have thus far supported the synonymization. When I discovered *E. pedicellatum* in western Labrador and in Quebec’s North Shore, I noticed that the specimens resembled the European specimens more than the ones from Atlantic Canada. A closer examination of European, Asian, and western North American specimens revealed that there appeared to be two forms of *E. pedicellatum* distributed along a latitudinal gradient. The southern form is mostly restricted to the island of Newfoundland and Nova Scotia while the northern form is mostly found in Alaska, Norway, Northern Quebec, and Gros Morne, and Labrador. Having many pictures of both forms as well as good voucher specimens from Canada, we are in a unique position to rigorously describe these two forms.

Here we test the hypothesis that two distinct forms or morpohotypes of E. pedicellatum exist This paper has two main objectives: 1. Describe the southern and northern forms from morphological, ecological, and biogeographical perspectives, 2. Assess the estimated size of the population of the boreal felt lichen for these two forms and the implications for its conservation status at a global, national, and subnational levels.

## Methods

### Examination of field pictures

At a coarse scale, we investigated ecological and morphological characteristics from photos taken in the field and images from herbarium specimens from North America, Europe, and Asia. Most photos were from Inaturalist and our own field photos from Canada. These were complemented by using a few photos available online from other sources. We also included the type specimen of *E. boreale* from Herbarium O, University of Oslo and included *P. pedicellata* from the Museum of Natural History of London. The ecological characteristics that we examined included, geographic region, the species of phorophyte, the microhabitat (branch or trunk) presence of the liverwort *Frullania*. The presence of *Lobarion* species adjacent to the boreal felt lichen (excluding *Parmeliella* and *Fuscopannaria*) was also noted from photos and field notes. “We also examined photos for differences in morphological characteristics, such as thallus colouration, amount of tomentum, and the appearance of the apothecia, including the presence of pruina (waxy or matte colour), a light or dark exciple margin, and the presence of noticeable hairs around the exciple margin.

### Detailed morphological description and measurements

In-depth examinations of morphological characteristics were also carried out using fresh and voucher specimens collected during field studies between 2019 to 2025, and included specimens from four populations, the Avalon Peninsula (n=7), Bay D’Espoir (n=15), Gros Morne National Park (n=4), and Quebec’s upper Northshore (n=12). Cross-sections of thalli and apothecia were prepared by hand using a razor blade. Samples were mounted in water and observed under a Leica S9i dissecting microscope and a Leica DMLB compound microscope. Images of thalli and apothecia were captured with a Galaxy S23 phone and the Leica S9i dissecting scope. Microscopic images were captured similarly through the eyepiece of the Leica DMLB.

Chemical spot tests were performed on specimens using paraphenylenediamine in ethyl alcohol (P), sodium hypochlorite (C), 10% potassium hydroxide (K), and Lugol’s iodine (I) following Brodo et al. (2001). The thalli of representative specimens were tested with K, C, and P. Cross sections of apothecia were tested with K followed by the addition of I and observed under light microscopy. The pruina on the surface of apothecia were tested with polarized light under the compound microscope using two small sheets of polarized film.

For each population, cross sections were made of apothecia to look at differences in the colouration of the exciple and to measure the length of hairs extending from the exciple. For consistency, the length and width of ascospores, specifically from mature, convex 4-5 apothecia were also measured using specimens from the Avalon Peninsula, Bay D’Espoir, Gros Morne (n=48).

Our specimens are housed at the Forest Ecology Lab and the Canadian Forest Service Herbarium (CDFN) at the Atlantic Forestry Centre in Corner Brook, NL.

### Statistical analyses

Ascospore dimensions and hair lengths from Quebec and Gros Morne specimens were grouped into the northern morphotype, and those from Hall’s Gullies and Bay D’Espoir were grouped into the southern morphotype.

Statistical analyses were performed using R. The normality of distribution of variables was checked using the Shapiro-Wilk’s test. Ascospore dimension data was not normally distributed so the Mann-Whitney U-test was used to test for significant differences in spore length, width, and length:width ratio (L/W). The hair length data was normally distributed so the Welch’s t-test was used to test for significant differences the length of exciple hairs

## Results

### Distribution ecology and morphology from field photos

The photos of the type specimens collected by Farlow in 1902 and Ahlner in 1941 show clear differences associated with the exciple of the apothecia which are consistent from our analysis of field photos (Fig 1).

**Fig. 1.**
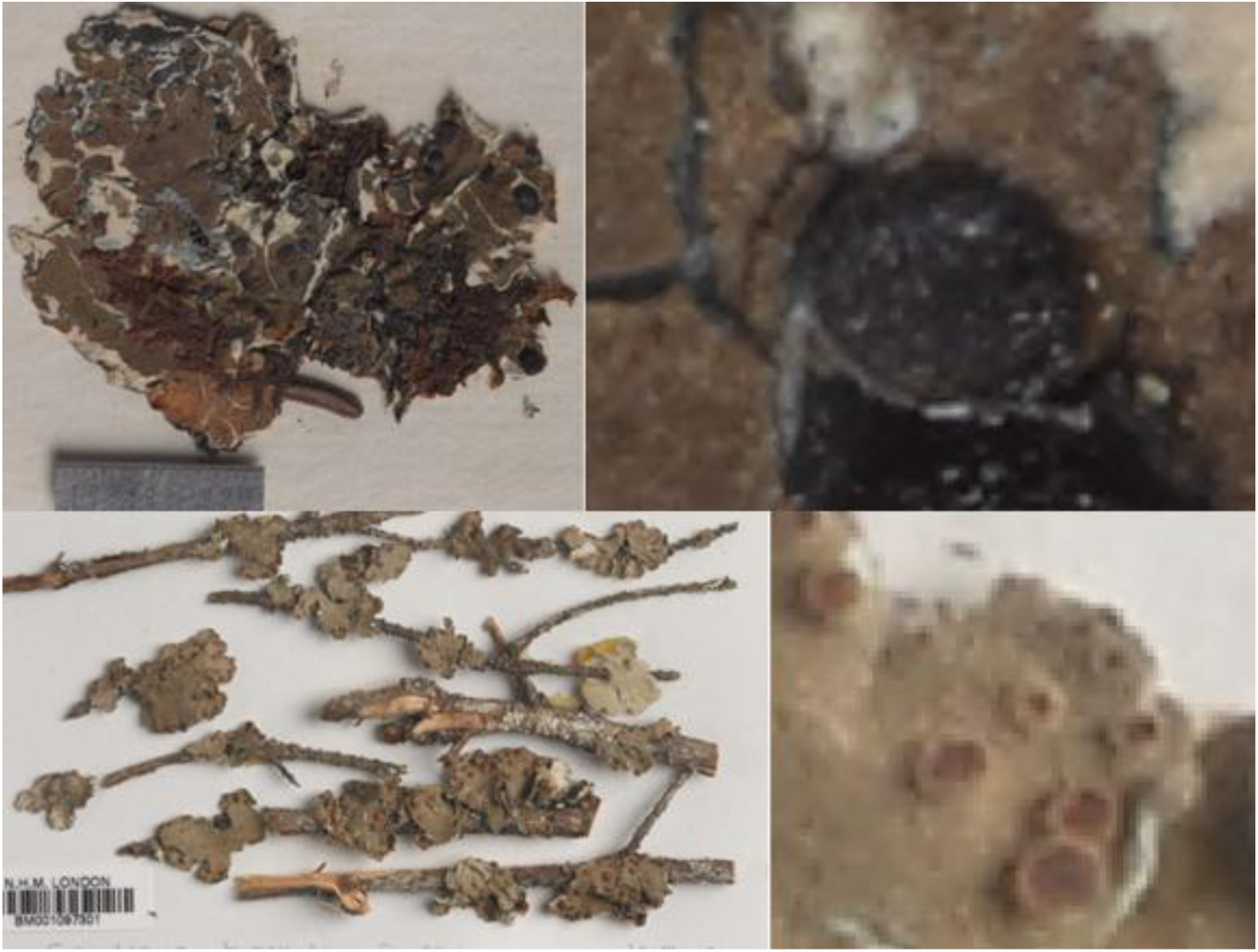
Photos of original voucher specimens of P. pedicellata described by Hue (NHM-London-BM001097302) (top) and *E. boreale* (NHM-London-BM001 097301) (bottom). On the right we added a magnified imaged showing the apothecia colour and exciples. These samples were examined by Jørgensen (1972) resulting in the synonymization of the two taxa into *E. pedicellatum*. These were respectively accessed at NHM-London on October 14 2026 at https://data.nhm.ac.uk/dataset/56e711e6-c847-4f99-915a-6894bb5c5dea/resource/05ff2255-c38a-40c9-b657-4ccb55ab2feb/record/3070908 and https://data.nhm.ac.uk/dataset/56e711e6-c847-4f99-915a-6894bb5c5dea/resource/05ff2255-c38a-40c9-b657-4ccb55ab2feb/record/3070906

The photos of field specimens and collected specimens show a clear pattern differentiating the two forms of *Erioderma pedicellatum* in terms of associated communities and phorophyte species. The southern form, except for 1 in 111 photos, was never seen growing near *Lobarion* lichens. Whereas most of the specimens of the northern form, except those from Quebec and Labrador, were regularly growing near a variety of *Lobarion* lichens. The southern form was almost always accompanied by the liverwort *Frullania asagrayana* (92% of photos) while this liverwort was never seen in association with the northern form except for Newfoundland. The southern form of the boreal felt lichen is predominantly found on balsam fir, *Abies balsamea*. On the Avalon peninsula and in Nova Scotia thalli were always found on balsam fir trunks while in Bay D’Espoir, approximately half were on trunks and half on branches. The northern form, throughout its range, is always located on the branches of conifers, never on the trunks. According to photos, the thalli from Quebec and Labrador were only on branches of balsam fir. However, we have records to show that it also occurs on *Picea mariana* and Anderson et al. (2022) reported a specimen on branches of *Picea glauca*. The host tree species were *Picea glauca* in Alaska and Gros Morne, *Picea abies* in Sweden and Norway, and *Picea jezoensis* in Kamchatka.

There were no clear morphological differences in the colouration of the thallus, in the amount of tomentum, or the shape of apothecia between the two forms in photos or in cross sections. Apothecia do seem somewhat more bulbous in the southern form and flatter in the northern form. But variability in colouration of apothecia and thalli, in the shape of apothecia, and in the amount of tomentum often existed within the same population, not just between populations. Variation seems strongly associated with thallus age, hydration, health and possibly other environmental conditions, such as light.

However, three morphological characteristics are consistently different between the two forms. The apothecia of the southern form regularly have a sheen, a waxy or frosted finish, suggesting the presence of pruina. The presence of pruina was rarely seen in the northern form. The exciple margin of apothecia in the southern form is always dark as opposed to the white or pale margin in the northern form. These differences are also noticeable on photos of the type specimens *Pannaria pedicellata* (Hue) collected by William Gilson Farlow in 1902 in New Brunswick (NHM-London-BM001097302), and the type specimen *Erioderma boreale* ahlner collected by Sten Ahlner in 1941(NHM-London-BM001 097301) (Fig. 1). Associated with the white margin in the northern form is the presence of a noticeable white fringe of hairs. These differences in the two forms are consistent throughout all the regions (Fig. 2), although not all photos are of a high enough resolution to clearly see these characteristics.

**Fig. 2.**
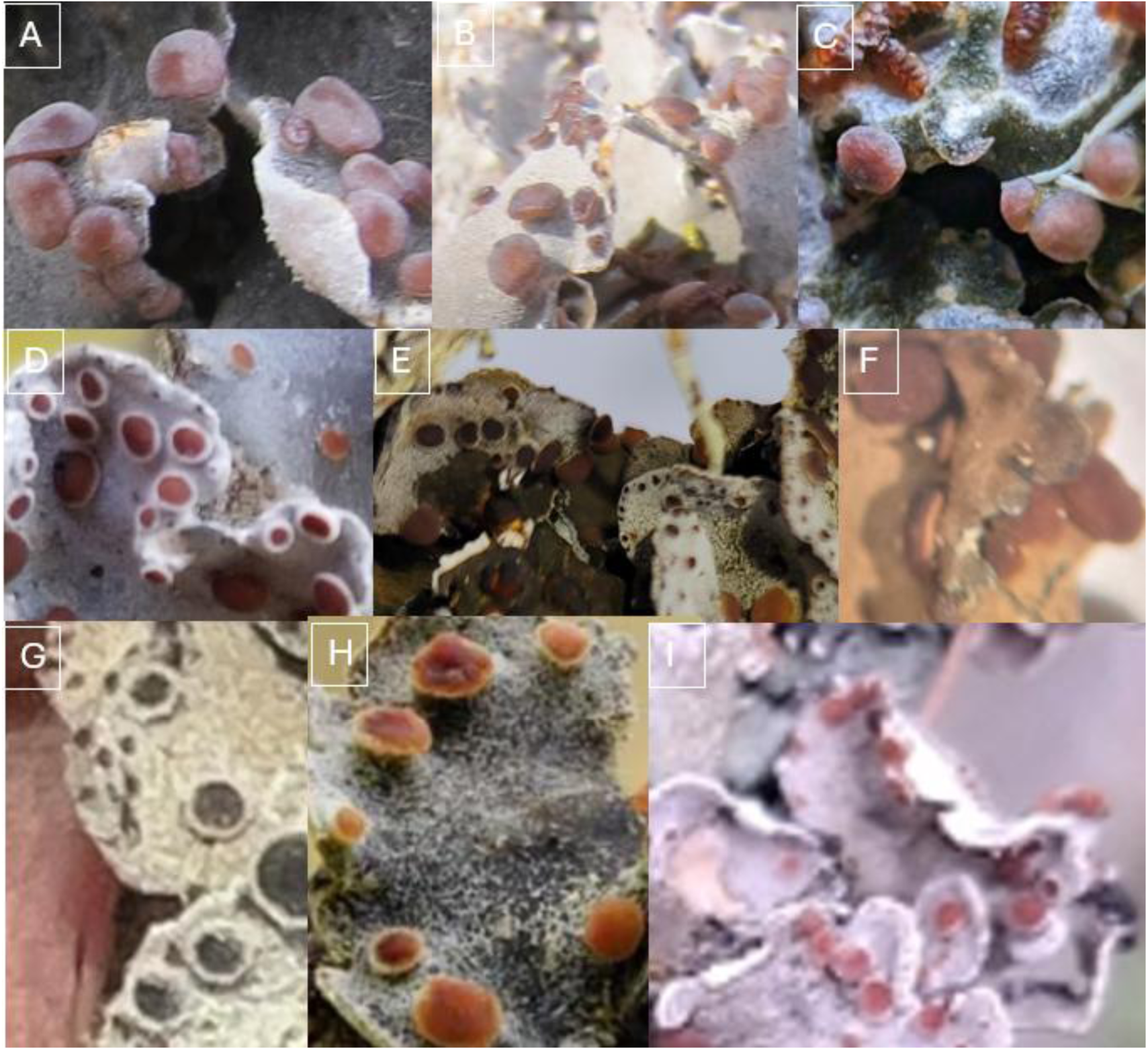
Photos comparing the boreal felt lichen from different locations. All photos are available online from Inaturalist unless indicated otherwise. Field photos of the southern morphotype (A) Halls Gullies from André Arsenault, INAT410065821 (B) Bay D’Espoir, from André Arsenault, INAT410060678, (C) Nova Scotia, from James Churchill, INAT365560414. Field photos of the northern morphotype (D) Quebec, from André Arsenault, INAT410067261, (E) Gros Morne, from André Arsenault INAT410068602 (F) Labrador, from André Arsenault INAT410067888 (herbarium specimen), (G) Alaska, from Preston Villumsen INAT344356316, (H) Norway, from Einar Timdal, artsdatabanken.no/arter/takson/47170, (I) Kamchatka from Violetta Dzizyurova, INAT410065821.

### Detailed morphology of the apothecia

Consistent with observations from field and specimen photos, microscopy revealed the presence of a dark exciple rim with relatively short hairs in the southern form (Fig. 3), while the northern form has apothecia with a light exciple rim with longer hairs (Fig. 4).

**Fig. 3.**
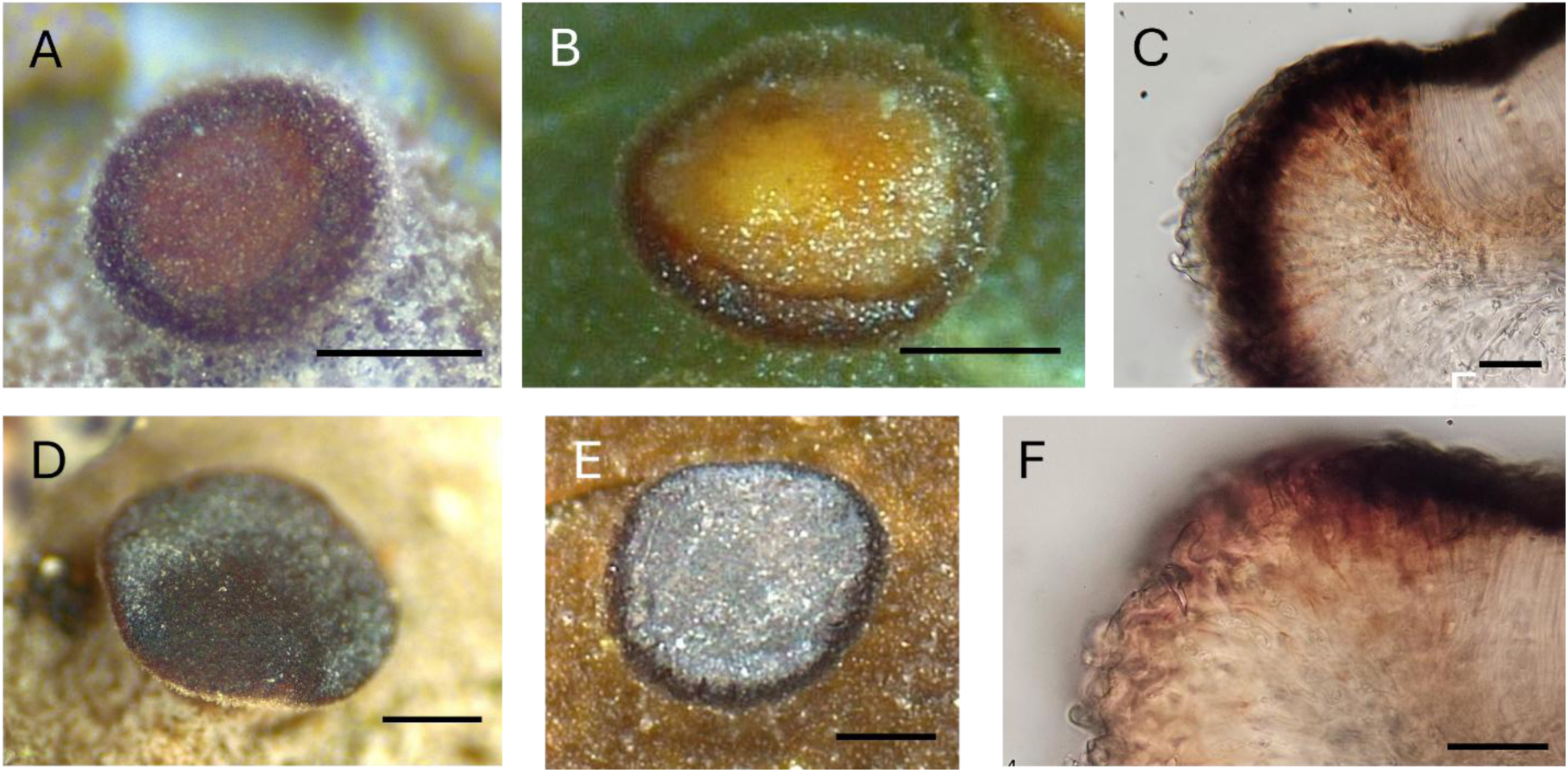
Flat to slightly convex apothecia with a dark rim, commonly seen in the southern morphotype. (A) Dry apothecium from the Avalon. (B) Wet apothecium from the Avalon. (C) Exciple of the apothecium from the Avalon, in water. (D) Dry apothecium from Bay D’Espoir. (E) Wet apothecium from Bay D’Espoir. (F) Exciple of an apothecium from Bay D’Espoir, in water. Scale bars: A, B, D, E = 0.25 mm, C & F = 25 µm.

**Fig. 4.**
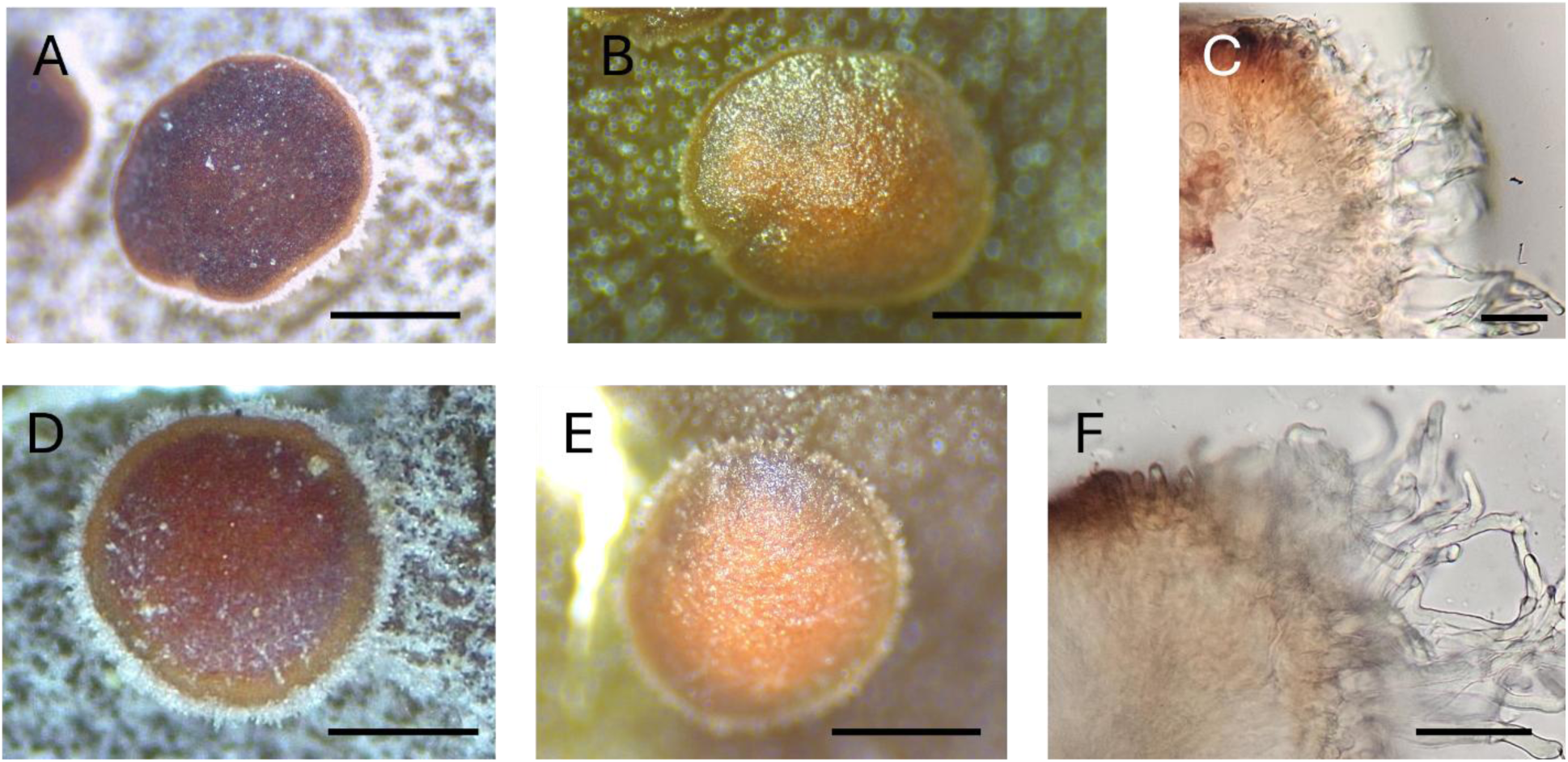
Flat to slightly convex apothecia with a white fringe of hairs, commonly seen in the northern morphotype. (A) Dry apothecium from Gros Morne. (B) Wet apothecium from Gros Morne. (C) Exciple of the apothecium from Gros Morne, in water. (D) Dry apothecium from Quebec. (E) Wet apothecium from Quebec. (F) Exciple of an apothecium from Quebec, in water. Scale bars: A, B, D, E = 0.25 mm, C & F = 25 µm.

Cross sections showed that apothecia on specimens from the Avalon and most specimens from Bay D’Espoir, the southern form, have a reddish-brown outer exciple (Fig. 3C & F). In specimens from the Quebec and Gros Morne the outer exciple is a hyaline to very light brown (Fig. 4C & F). The darker exciple gives apothecia of the southern morph a visibly darker rim that is especially apparent when apothecia are wet (Fig. 3).

In both forms apothecia become more convex and bulbous with age and the hairs are no longer visible from above in the northern form. The colour of apothecia often darkens with age.

The development of *Erioderma pedicellatum* is shown in Figs. 5 and 6. Pycnidia and primordial apothecia are found on the margins of young lobes of both forms. Juvenile and early developmental stages of apothecia have a raised proper margin with a rim covered in light hairs. In the southern form, by the time apothecia reach the flat to slightly convex stage of development they no longer have visible hairs on the margins (Fig. 5A & D). Apothecia of the northern form, on the other hand, retain the white fringe of hairs in the flat and into the slightly convex stage of development (Fig. 6A & D). In both forms apothecia become more convex and bulbous with age and the hairs are no longer visible from above in the northern form. The colour of apothecia often darkens with age.

**Fig. 5.**
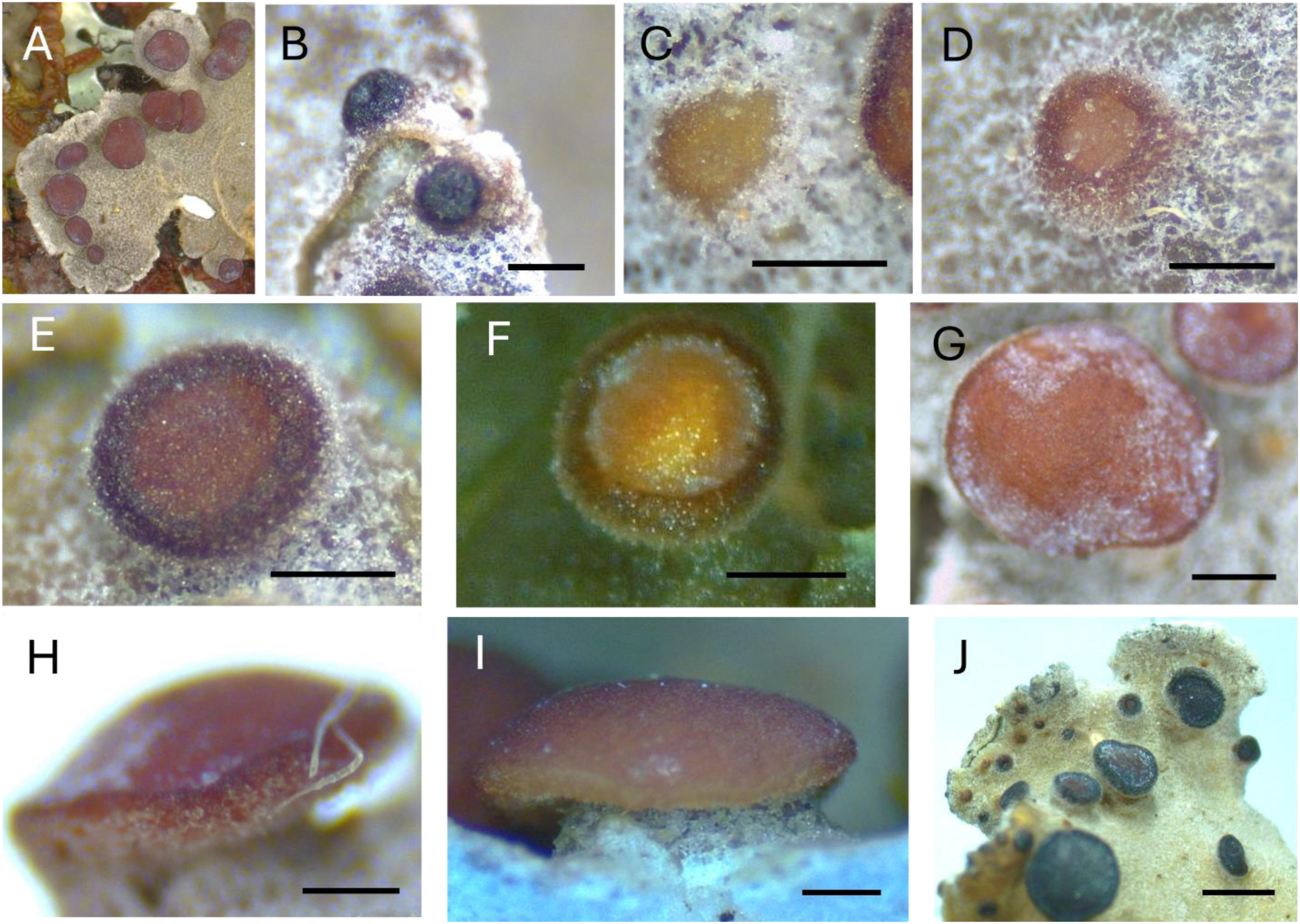
Images showing reproductive structures and development of apothecia in the southern form of *Erioderma pedicellatum*. (A) Thallus lobe with apothecia. (B) Mature pycnidia on the margin of a young lobe. (C) Primordial apothecia on a young lobe margin. (D) Developing apothecia, still immersed in cortex. (E) Young apothecia raised above the cortex. (F) Young apothecia, wet. (G) Flat apothecia with pruina. (H) Convex apothecia. (I) Bulbous apothecia. (J) Diversity seen in thalli, this one has very dark apothecia. Scale bars: B: 0.1 mm, C-I: 0.25 mm, J= 1 mm.

**Fig. 6.**
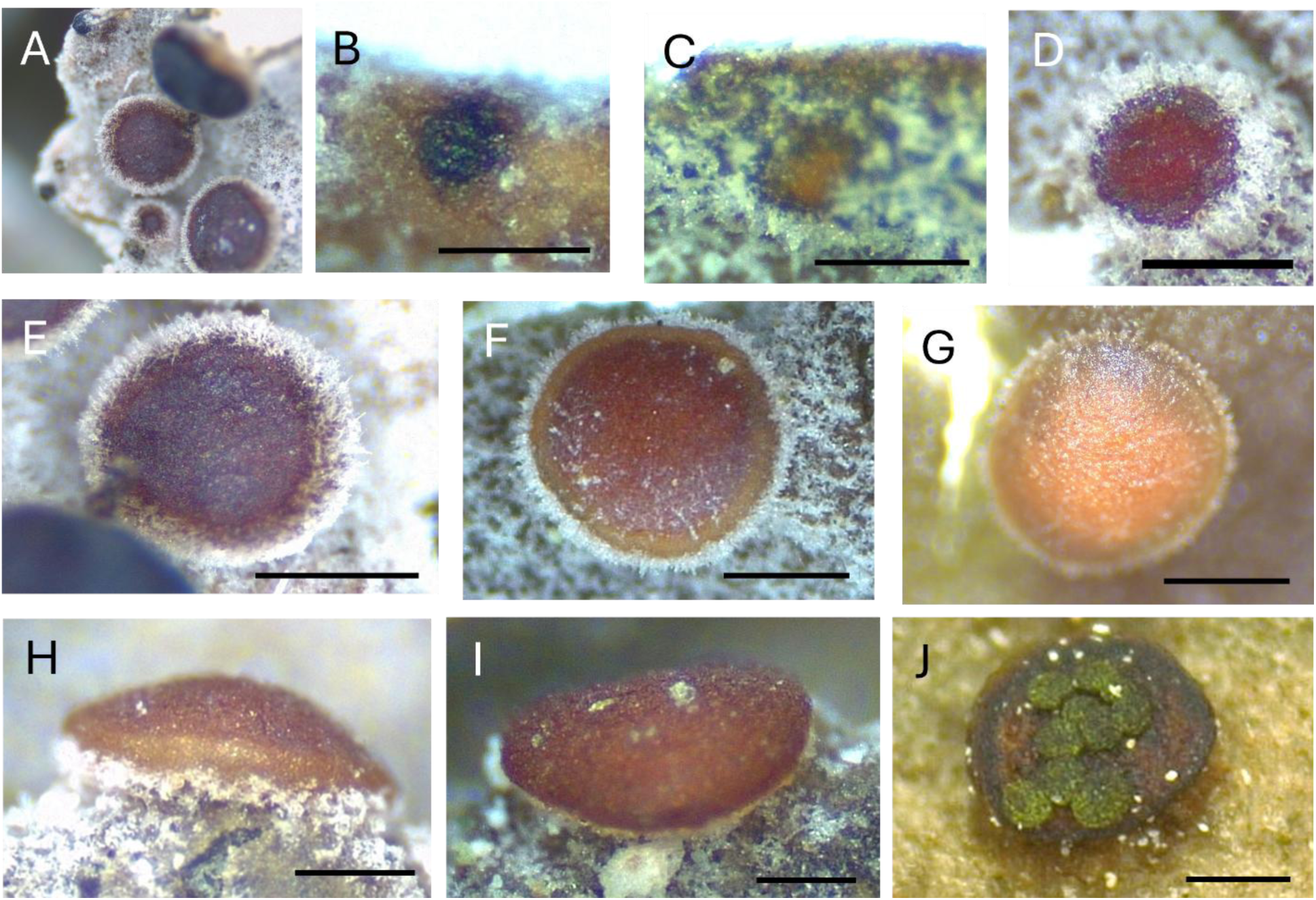
Images showing reproductive structures and development of apothecia in the northern form of *Erioderma pedicellatum*. (A) Young lobe with apothecia. (B) Mature pycnidia on a lobe margin. (C) Primordial apothecia on a young lobe margin. (D) Developing apothecia, still immersed in cortex. (E) Young apothecia raised above the cortex. (F) Flat apothecia. (G) Flat apothecia, wet. (H) Convex apothecia. (I) Bulbous apothecia. (J) Apothecia colonized by *Abrothallus nitassinan*. Scale bars: B: 0.1 mm, C-J: 0.25 mm.

### Microscopic measurements

We compared the length of exciple hairs on the flat to slightly convex apothecia from the northern form (Quebec) and the southern form (Halls Gullies and Bay D’Espoir combined) and found that the hairs on apothecia of the northern form were significantly longer than those on apothecia of the southern form (Fig. 7, Table 2).

**Fig. 7.**
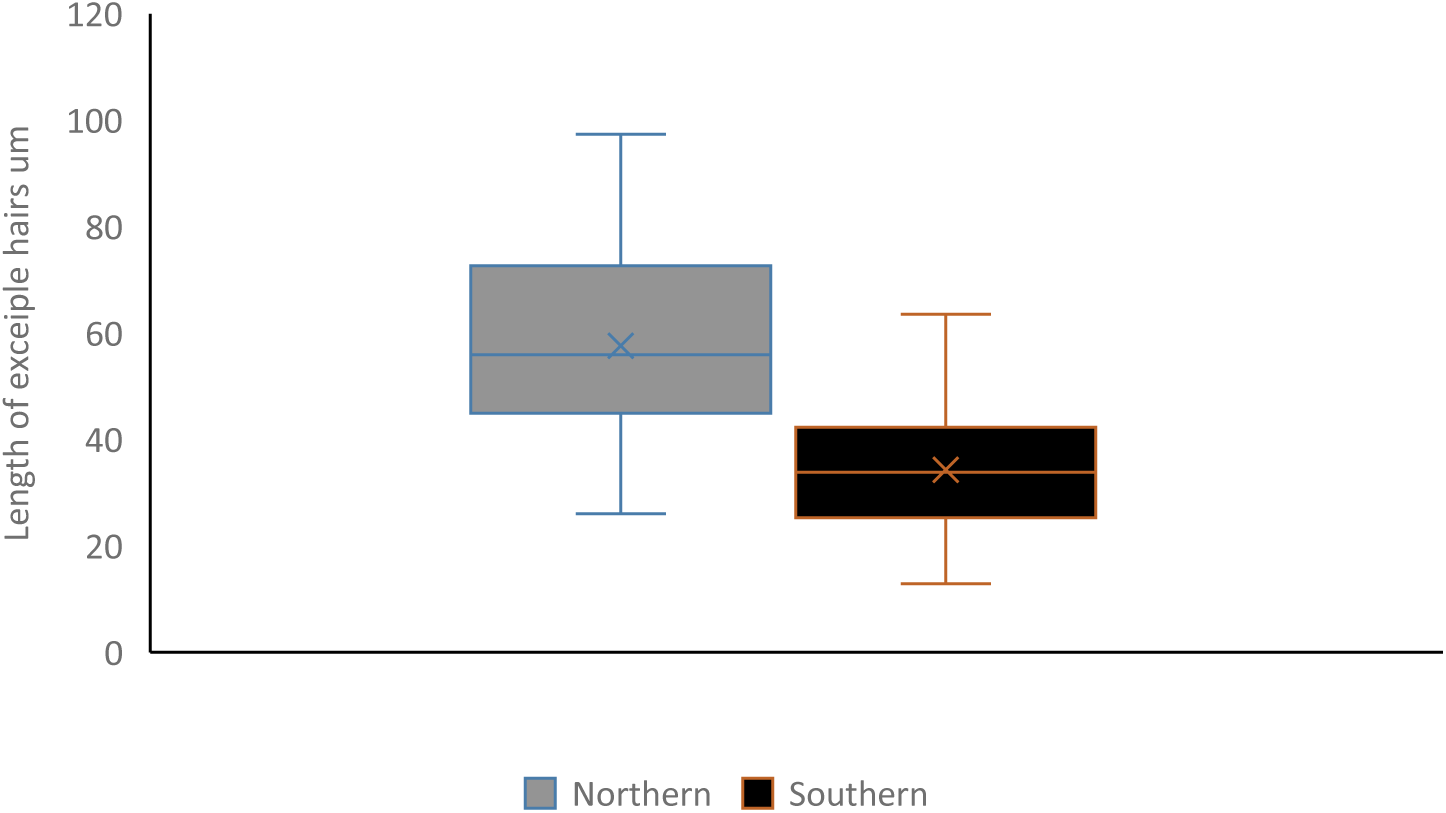
Length of exciple hairs for the northern form (grey) and southern form (black) of Erioderma pedicellatum, (p-value = 2.9e-07).

**Table 2.** Comparison of the ascospore dimensions and the length of exciple hairs in the two morphotypes. P-value results are from the Mann-Whitney U-test for ascospore dimensions and from a Welch’s two sample t-test for hair length.

| Morphotype | Spore width (µm) |  | Spore length (µm) |  | Spore L/W ratio |  | Exciple hair length (µm) |  |
| --- | --- | --- | --- | --- | --- | --- | --- | --- |
|  | Mean | SE | Mean | SE | Mean | SE | Mean | SE |
| Northern | 8.48 | 0.085 | 14.06 | 0.128 | 1.67 | 0.019 | 57.63 | 3.38 |
| Southern | 7.58 | 0.081 | 12.93 | 0.14 | 1.72 | 0.025 | 34.32 | 1.89 |
| p-value | 2.50E-11 |  | 3.04E-08 |  | 0.3529 |  | 2.91E-07 |  |

We also compared the dimensions of ascospores from the two morphotypes. The data was not normally distributed so a Mann-Whitney U-test was used to compare the ascospore dimensions (Table 2). The length and width of ascospores of the northern form ((11)-14.1-(17.5) x (6.7)-8.5-(10.5) µm) were significantly larger than those of the southern form (10)-12.9-(16.5) x (6)-7.6-(9.5) µm). In terms of spore shape, the length to width (L/W) ratio was not significantly different, the ratio range was 1.2-2.1 for the northern form ascospores and 1.3-2.5 for the southern form ascospores.

### Evidence of pruina and chemistry

Consistent with observations from photos of field and collection specimens, closer observation of apothecia on specimens from Hall’s Gullies, Bay D’Espoir, Gros Morne, and Quebec indicate that the southern form often has apothecia with a sheen, a waxy or frosty pruina, on the apothecial disk (Fig. 3D & E). This pruina is rarely seen on apothecia of the northern form. The pruina tested negative under polarized light but tested P+ orange, indicating that the waxy or frosted appearance is most likely the result of secondary compounds rather than true pruina.

The chemistry of both forms was examined using specimens from the Avalon and Bay D”Espoir region of the Island of Newfoundland for the southern form and specimens from Quebec’s North Shore for the northern form Table 3. The chemistry is generally the same between the two forms with two exceptions, the exciple of the apothecia in the southern form turns from a reddish brown to darker brown with the addition of K while the exciple of the northern form is K-, and the southern form usually has apothecia with P+ orange pruina or secondary compounds, while the northern form does not.

**Table 3.** Spot test chemistry of the northern and southern morphotype of *Erioderma pedicellatum*.

| Form | Pruina | Thallus | Medulla | Excipule rim and apothecia | Hymenium |
| --- | --- | --- | --- | --- | --- |
| Southern | K-, C-, P+, UV- | K+ brown, C- | P+O | K+ Brown darker | I+blue |
| Northern | None | K+ brown, C- | P+O | K- | I+blue |

## Discussion

### Morphological variation of the boreal felt lichen

Our field work on the boreal felt lichen and the opportunities to observe and collect fresh material from multiple sites on the island of Newfoundland, western Labrador, and Quebec’s north shore, along with access to good online field photos from Nova Scotia, Alaska, Norway, Sweden, and Kamchatka and good photos of herbarium specimens from the Museum of Natural History of London, Herbarium S of Swedish Museum of Natural History and Herbarium O of the Natural History Museum at the University of Oslo, placed us in an advantageous position to be able to recognize, provide evidence for and describe two distinct morphotypes of the boreal felt lichen in the northern hemisphere. Table 4 compares key characteristics of the original descriptions of *Pannaria pedicellata* Hue by Hue, *Erioderma boreale* Ahlner by Ahlner, and their synonymization into *E. pedicellatum* Hue Jørg. by Jørgensen, along with our descriptions of the southern morphotype from Newfoundland and the northern morphotype from Quebec and Gros Morne National Park. The original description of the lichen as *P. pedicellata* Hue by Hue seems mostly consistent with our description of the southern form. He mentions that the exciple of the apothecia have a reddish-brown colour. Similarly, the original description by Ahlner of *E. boreale* also closely matches our description of the northern form. He describes the exciple colour as yellow to tawny and mentions that the rim is covered in a short wool. Both did seem to miss some features and although Jørgensen noticed differences between the specimens from New Brunswick and Sweden, he considered them insignificant. These discrepancies were likely related to the quality and quantity of specimens available to them at the time. “Our work emphasizes the importance of using a range of live specimens and taking into account their ecological context when describing lichens.

**Table 4.**
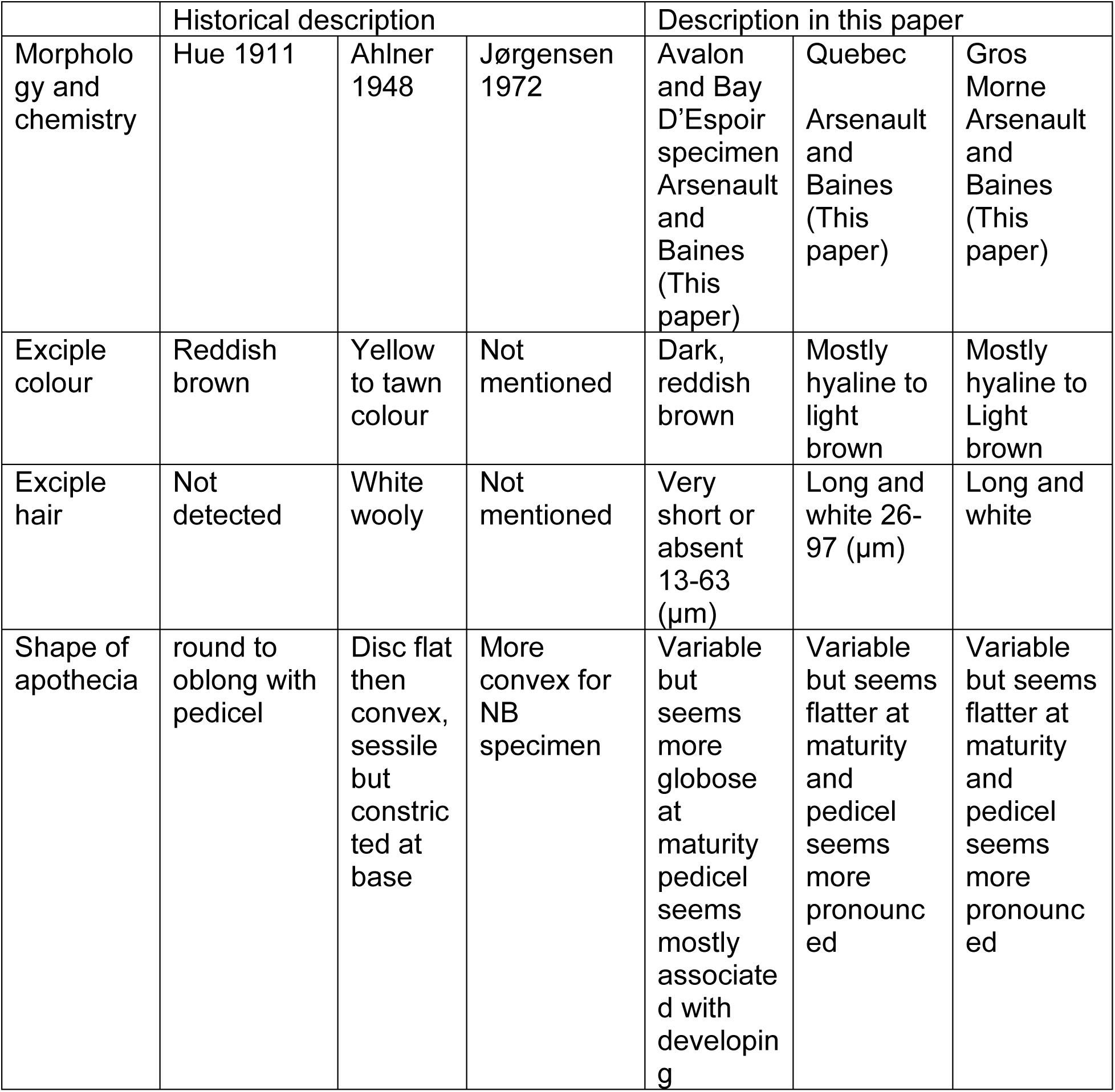
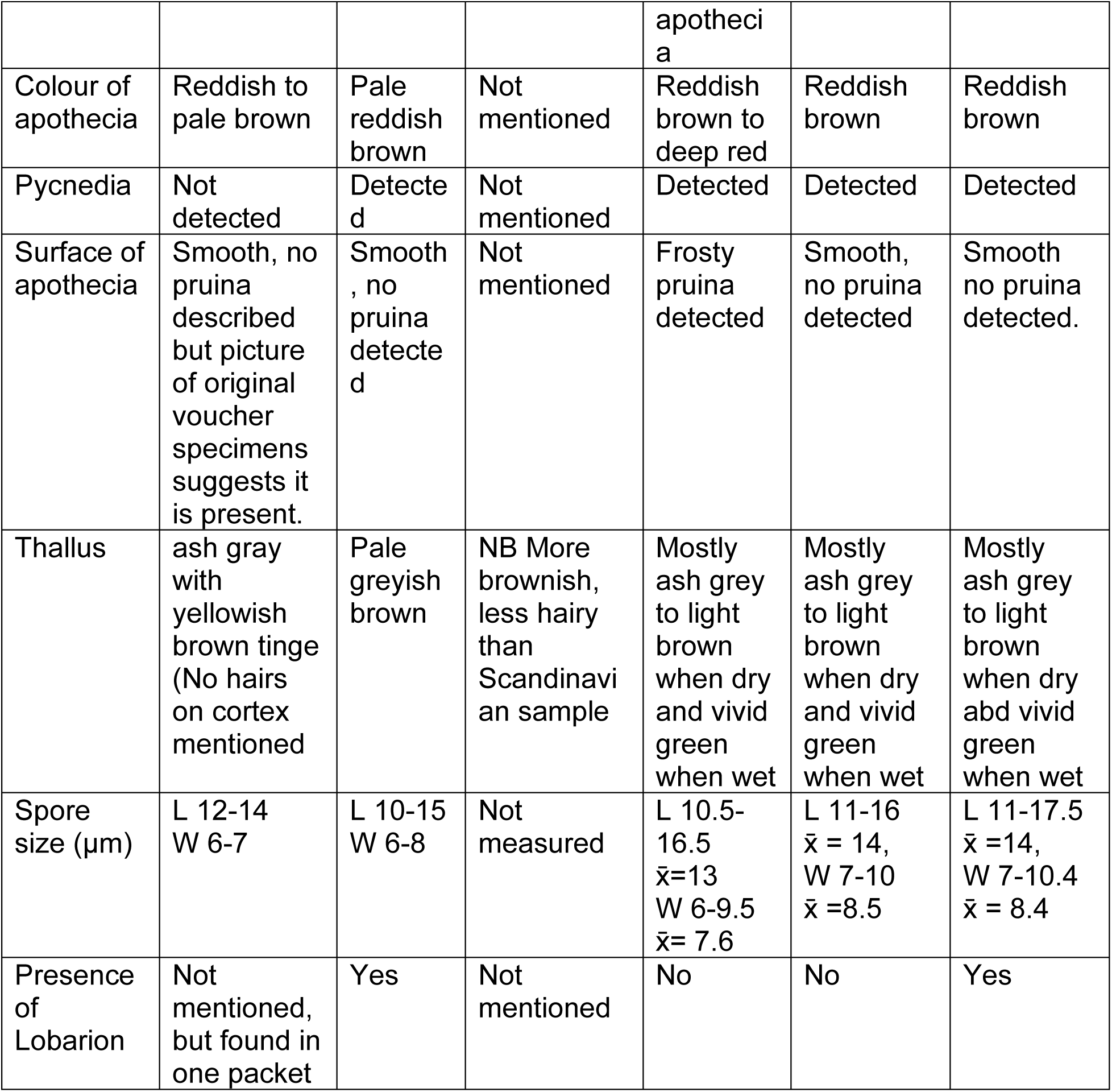
A comparison of descriptive traits for the boreal felt lichen using published description and our analysis of the two morphotypes we describe in this paper

### Distribution ecology of the two forms of the boreal felt lichen

#### Phoprophyte Specificity

Although there are a few exceptions, the southern form is mostly found on the trunks of *Abies balsamea* and in some cases on branches as well, especially on the south coast of Newfoundland. This contrasts with the northern form that lives mostly on the branches of Picea, except for sites in Quebec and Labrador where it was found growing in abundance on the branches of *Abies* as well. This could suggest that the relationship between the northern form of the boreal felt lichen and the host conifer species is mostly coincidental and it is more strongly influenced by the tree species composition of the stand it which it lives (Holien et al. 1995). However, one striking observation that counters this theory is seen in the forests of Gros Morne National Park in northwestern Newfoundland, where *E. pedicellatum* has been observed only on branches of *Picea*, not *Abies*, despite the occurrence of fir trees in close proximity to *Picea glauca* occupied by the boreal felt lichen. This combined with its sympatric occurrence with cyanolichens belonging to the *Lobarion* community suggest a requirement for unique bark characteristics.

#### Community ecology and possible species interactions

The presence of the *Lobarion* adjacent to the boreal felt lichen on the same branches is a characteristic of the northern form and mostly absent for the southern form. The *Lobarion* is often associated with forests of high humidity and stability (i.e. old-growth), and circum-neutral to basic substrates. It is thus possible that the northern form requires higher pH as suggested by Nillsson et al. (2022). It is difficult to know at this point whether the higher bark pH in Norway is a habitat requirement of the boreal felt lichen or just coincidental. If the former, cyanolichens of the *Lobarion* co-habiting on the same branches could facilitate the establishment of the boreal felt lichen, as the cyanolichens of the *Lobarion* are known to enrich bark pH (Arsenault and Baines 2026), or the co-existence could be just by chance and related to a high moisture environment.

Although the morphology of the northern form seems identical across its range in North America, Europe, and Asia, the community ecology patterns for the lichen in Quebec’s Manicouagan region and western Labrador deviate from the other subpopulations characterized by the absence of the *Lobarion*. This could suggest that the association with the *Lobarion* and specific conifer species are stochastic or that there are there are some other variations involved in the population structure across the species range.

The association of the southern form with the liverwort *Frullania asagrayana* Mont. was found in over 92% of the field pictures. This is not a new observation (Maas 1980, 1983 Scheidegger 1998). In Newfoundland, there is evidence of a photobiont mediated lichen guild (COSEWIC 2002, Cornejo and Scheidegger 2016, Cornejo et al. 2016) whereby there are possible exchanges of photobiont between *Coccocarpia palmicola*, *Erioderma pedicellatum* and the liverwort and possible facilitation of lichenization by the latter in its watersacs. The northern form does not grow on *Frullania* which implies a different process is involved in lichenization. In addition, Coccocarpia palmicola which predominantly occurs with the southern form, co-occurs with the Gros Morne subpopulation and one of Quebec’s subpopulations (Arsenault and Baines 2026).

There are possibly other biotic relationships involving the boreal felt lichen that could be explored but one last one worthy of mention involves lichenicolous fungi. A new species of lichenolous fungi, Abrothallus nitassinan Arsenault, P. Baines & Suija, was found to colonize thalli of Erioderma pedicellatum in the northern form of Quebec and the southern form in Bay D’Espoir (Suija et al. 2024). Although sometimes these fungi are described as parasites of lichen, their ecological roles and inter-species relationships are poorly understood (Lubek et al. 2019). In both the Quebec and Newfoundland examples, branches where mature thalli of the lichen had been colonized and severely affected by these fungi by these fungi had very abundant juvenile thalli. It is therefore possible that these fungi facilitate some of the processes associated with the ejection, and germination of spores, or alternatively environmental conditions are both optimal for the lichen and the lichenicolous fungi.

Based on our comparison of species description and consistent patterns of distribution of the northern versus southern forms, except for Quebec and Labrador, we propose that their respective differences should not be simply dismissed as the result of ecological conditions sensu Jørgensen (1972). New genomic analyses will be required to untangle the population structure of the boreal felt lichen.

### Current status of the population of the boreal felt lichen

The global assessment of the boreal felt lichen by the IUCN is over twenty years old (Scheidegger 2003) and was based on an enumeration of 5,074 only found in Atlantic Canada. Furthermore, the historical subpopulations of New Brunswick, Sweden and Norway were considered extinct. Much new information has surfaced since then (Figure 8), including new populations found in Asia (Stepanchikova & Himelbrant 2012), in western North America (Nelson et al. 2009) and in the boreal mainland of Canada (Anderson et al. 2022, Arsenault 2022) as well as new subpopulations in Norway reviving this lichen from several presumed extinctions (Jørgensen 1990, Sheidegger 2003). Recent assessments of the boreal felt lichen at national scales include updated estimated or inferred population sizes of 15,500-20,500 for Canada (COSEWIC 2014), 100,000 for the USA (Stehn et al. 2013), 1500 for Norway, and more than 100,000 for Russia (Tagirdzhanova et al. 2019) This represents a global estimate of over 217,000 mature individuals which represent a 42 fold increase since the last assessment and a major increase in the extent of occurrence of this lichen.

**Figure 8.**
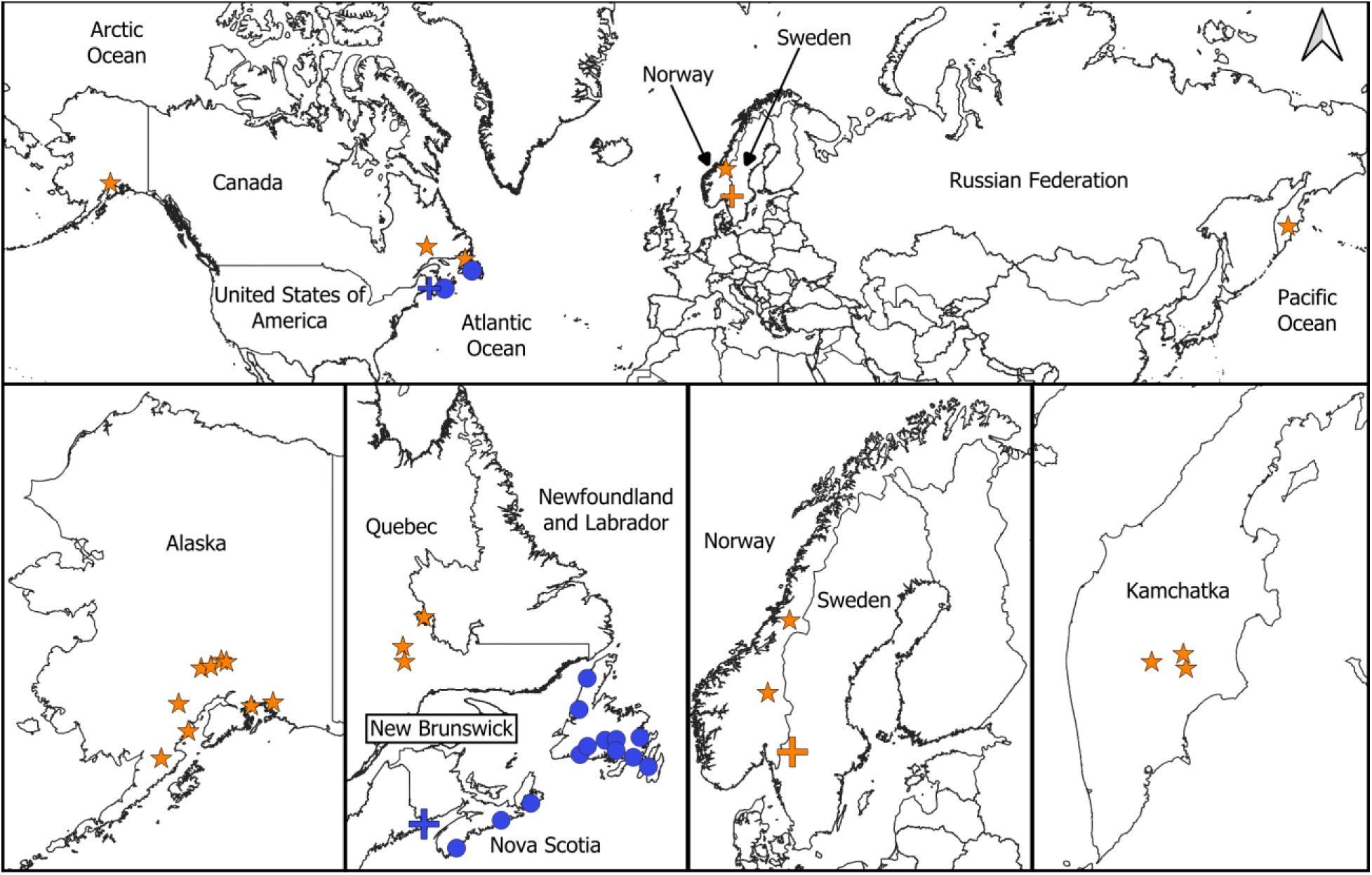
The distribution of the boreal felt lichen as of October 16, 2026. The northern morphotype locations are shown as orange stars while the southern morphotype locations are shown as blue dots. The extinct subpopulations are shown as +.

Although, new assessments of this lichen are urgently needed at global, national and sub-national scales, several factors make this complicated. For example, the original designatable unit (DU) structure in Canada which resulted in an assessment of an Atlantic and a boreal population is no longer supported from data presented here. Furthermore, one of the main reasons for creating these two populations was due to a higher level of threat in the Atlantic population (COSEWIC 2002) which is not an acceptable criterion for DU assessment (Green 2005). Other complications include recent mortality of host trees in Alaska and Newfoundland resulting from insect outbreaks (Perez and Stehn 2025, Arsenault personal observation), and severe declines of mature individuals in Nova Scotia resulting from herbivory from introduced gastropods (Cameron and Toms 2016). In some cases, like the forests of Gros Morne National Park, detection may be a challenge as the lichen occurs often at height 3 metres above the ground. Furthermore, in the cases where the lichen grows on branches, it can be so abundant as to making accurate assessments of the number of thalli very difficult. Therefore, using a single occupied tree as a single functional individual for arboreal lichen is likely to be a more robust approach in lichen assessments (Yahr et a. 2019) and this approach has now been adopted by COSEWIC. In addition, there are vast areas of un-surveyed potential suitable habitat which could significantly increase the estimated population. This combined with the fact that the boreal felt lichen follows meta-population dynamics that is not well captured by current assessments will be present a significant challenge in any future assessment. New field research and monitoring will be required to do robust assessments. In addition, in forests where the thalli grow on branches of conifers, often in great abundance, it is impractical to attempt to estimate the number of thalli for conservation assessments. Therefore, using a single occupied tree as a single functional individual for arboreal lichens (Yahr et al. 2019) is a more realistic approach and is now being adopted by COSEWIC in Canada.

Our understanding of the biology and ecology of lichens has increased significantly in the last few decades (Brodo et al. 2001, Goward and Arsenault 2001, Scheidegger and Werth 2009, Gauslaa, 2014, Spribille et al. 2016, Ellis 2019, Lücking, and Spribille 2024, Goward 2024). However, there are still many unanswered fundamental questions which if resolved, could “enlichen” our assessment of conservation status and threats and assist in developing innovative mitigating approaches for species conservation. Factors involved in lichenization, dispersal, sexual reproduction, generation time, establishment and growth, and relationships with co-occurring taxa in different taxonomic groups, like lichenicolous fungi, represent only a few of these very important information gaps.

## Acknowledgements

We thank Nick Lake who has provided many constructive comments on previous drafts and has created Fig.1. We also thank Miawpukek First Nation natural resource managers Gregory Jeddore, and Angelina Francis and the land guardians, Andy Joe, Raymond Jeddore, David Jeddore, Pat Jeddore, Roger Macdonald, and Greg Benoit, and many others for much insight into the distribution of the boreal felt lichen and the land and assisting in training and surveys of rare lichens in Bay D’Espoir, Terra Nova National Park, and Gros Morne National Park. We thank our collaborators from the Pessamit Innu to work in the Nitassinan with them on the caribou project. We thank Parks Canada, especially John Gosse, Darroch Whittaker and Lena Dietz Chiasson for assisting with rare lichen surveys in Terra Nova and Gros Morne National Parks. The work we did in Gros Morne and Terra Nova National Parks were under a research permit TNP-2019-32739. Finally, we thank all our colleagues who worked directly with us in the field, feu Bill Clarke, Claudia Hanel, Janet Feltham, Robert LeBlanc, Candice Power, Randy Skinner, Vanessa Hussey, Carlos Pasiche Lisboa, Katherine Flores, Adam Green, Chris Ratcliff, and many others including scientists who contributed valuable knowledge on the biology of the boreal felt lichen since 1902. We thank Daniel Stanton, Toby Spribille, and Trevor Goward for email exchanges that helped determine that the location of the voucher listed as Summit Lake, British Columbia was likely a mistake. We thank the many contributors to online databases such as Inaturalist, GBIF, and Consortium of North American herbaria. Over the last 16 years funding has been provided by several agencies including the Canadian Forest Service Research Program, Natural Resources Canada Assistant Deputy Minister Innovation Fund, Canadian Forest Service Assistant Deputy Minister Innovation Fund, NSERC, Newfoundand and Labrador Forest Service Centre for Science and Evolution, as well as Corner Brook Pulp and Paper.

